# Osseointegration of functionally-graded Ti6Al4V porous implants: Histology of the pore network

**DOI:** 10.1101/2023.01.05.521963

**Authors:** Joseph Deering, Dalia Mahmoud, Elyse Rier, Yujing Lin, Anna Cecilia do Nascimento Pereira, Silvia Titotto, Qiyin Fang, Gregory R. Wohl, Feilong Deng, Kathryn Grandfield, Mohamed A. Elbestawi, Jianyu Chen

## Abstract

The additive manufacturing of titanium into porous geometries offers a means to generate low-stiffness endosseous implants with a greater surface area to improve osseointegration. In order to optimize pore size in the scaffolds, it is important to first understand the timeline of osseointegration in pre-clinical models. In this work, selective laser melting was used to produce gyroid-based scaffolds with a uniform pore size of 300 μm or functionally-graded pore size from 600 μm to 300 μm before implantation in New Zealand white rabbit tibiae for 4 and 12 weeks. Initial *in vitro* assessment with Saos-2 cells showed favourable cell proliferation at pore sizes of 300 and 600 μm. At four weeks, histological observations indicated some residual inflammation alongside neovessel infiltration into the scaffold interior and some early apposition of mineralized bone tissue. At twelve weeks, both scaffolds were filled with a mixture of adipocyte-rich marrow, micro-capillaries, and mineralized bone tissue. X-ray microcomputed tomography showed a higher bone volume fraction (BV/TV) and percentage of bone-implant contact (BIC) in the implants with 300 μm pores than in the functionally-graded specimens, indicating that these smaller pore sizes may be favourable for osseointegration in leporine bone.

**Graphical Abstract:** 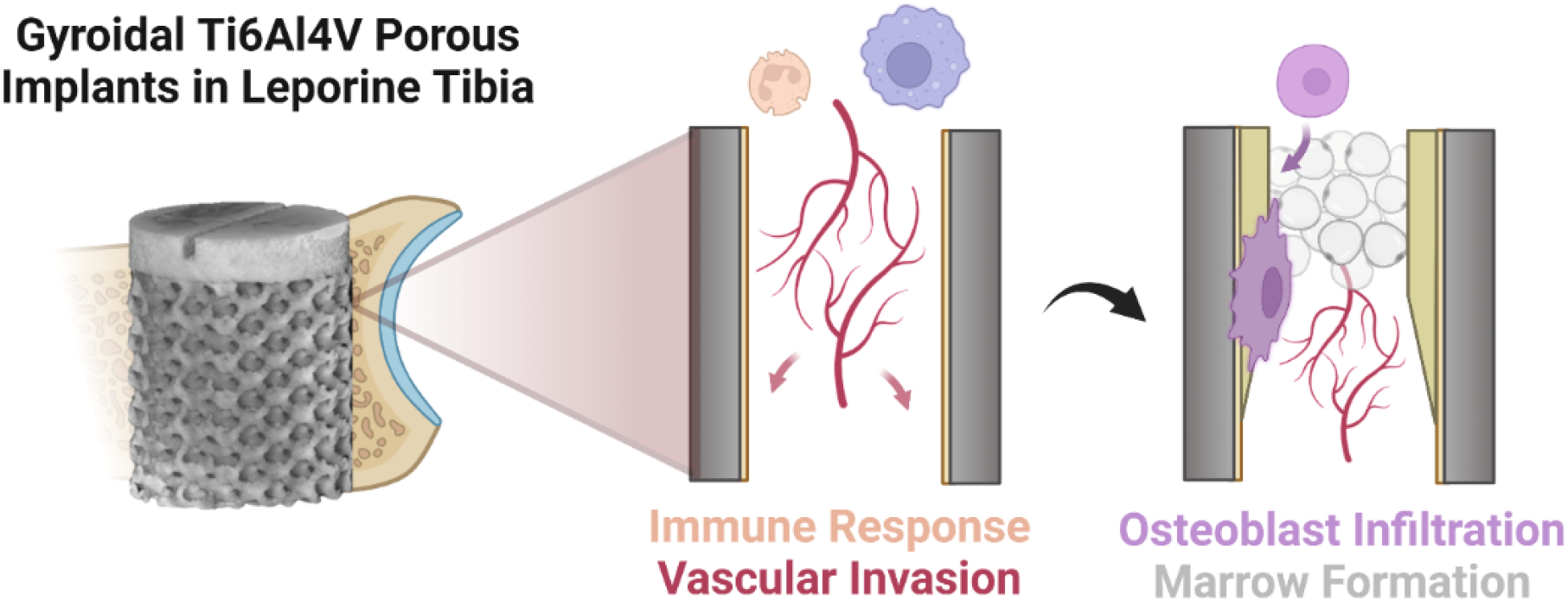

## Introduction

Central to the success of dental and orthopaedic implants is the development of a structural and functional connection at the bone-implant interface [1]. Careful control over material biocompatibility, surgical technique, and macroscale implant design are among the core parameters for governing the early stages of osseointegration – especially important as failure is most common within the first year of implantation [2]. These design strategies [3,4] have been developed to include elements of isotropy [5,6] and biomimetics to tune the mechanical characteristics [7,8] of tissue scaffolds, with renewed interest in optimizing scaffolds for osseointegration. In particular, the advancement in laser powder bed fusion (L-PBF) has allowed for the fabrication and geometric control of metallic lattice structures for use in bone tissue engineering applications [9–11]. These lattice structures utilize interconnected pore structures to provide additional surface area for bone ingrowth [12] and their use aims to improve the secondary stability of dental and orthopaedic prostheses. Moreover, the porosity reduces the overall stiffness of the implant and potentially helps to mitigate bone resorption associated with stress shielding phenomena as suggested by numerical simulations with a low-stiffness implant stem [13].

In general, additively manufactured lattice structures can be categorized by their repeating topology into a dichotomy of computer-aided drawings (strut-based) or triply periodic minimal surfaces (TPMS) [14,15]. Gyroid is one subset of TPMS lattice family, with the possibility to mimic the mechanical properties of bone by tuning the relative density of the scaffold [16]. The main advantage of gyroid structures is that their architecture facilitates cell migration, in part due to their infinitely smooth architecture and relatively high fluid permeability [17]. TPMS structures also demonstrate high surface area to volume ratio, high energy absorption [18], and high specific strength/stiffness [19]. Recent research has been directed to investigate the capability of functionally-graded (FG) TPMS designs in terms of pore volume fraction and morphology [20–22]. While the optimal pore size for each pre-clinical animal model is still disputed, a benchmark for adequate pore size ranges between 300-500 μm for osseointegration to occur and porosity values in the implant ranging from 50-70% appear to be most abundant for porous implants [11]. Numerous experimental and numerical studies of functionally-graded TPMS scaffolds have shown they are suitable for bone tissue engineering based on their mechanical properties [23–25], but there is a distinct research gap relating to the osseointegration of additively manufactured implants with a changing pore size throughout their interior.

Despite the design and *in vitro* characteristics of TPMS structures having been reviewed elsewhere [26], these forms of experimentation are generally limited compared to *in vivo* studies [27]. In general, the *in vivo* wound healing response in the peri-implant environment follows distinct stages of clotting and inflammatory response, sprouting of new blood vessels and extracellular matrix deposition, and formation of mineralized bone tissue around classical implants [28]. However, there is a distinct lack of in-depth histological probes describing whether these biological processes occur through the interior of porous metals during the healing timeline – especially in implants with TPMS geometries.

In this work, we assessed the *in vitro* cytocompatibility and *in vivo* osseointegration within the interior of uniform and functionally-graded Ti6Al4V gyroid structures. Using Saos-2 metabolic activity measurements, X-ray microcomputed tomography of newly formed tissue in the leporine tibia, and hematoxylin/eosin staining of histological sections, we describe common themes in bone tissue nucleation and adaptation over twelve weeks within the sheltered porous interior of metallic implant materials.

## Methods

### Scaffold Design and Fabrication

TPMS Ti6Al4V gyroid parts were fabricated by selective laser melting additive manufacturing. The Renishaw additive manufacturing machine was equipped with a 200 W pulsed laser, creating a series of overlapping exposures to assemble parts one layer at a time using an exposure time of 40 μs, point distance of 60 μm, layer thickness of 30 μm, and hatch spacing of 65 μm [29]. The surface of gyroids was modelled using Equation 1, where *a* defines the unit cell size, *t* defines the volume fraction (relative density) of the gyroids, and *x/y/z* represents the spatial coordinates within the scaffold interior.

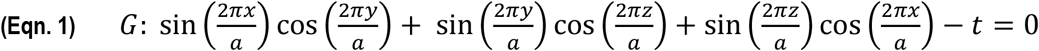

A MATLAB® code was used to generate the surface of the gyroids, both with functional grading and without (i.e., uniformity) in the pore structure. Uniform gyroid structures with an approximate pore size of 300 μm (herein denoted as G300), uniform pore size of 600 μm (G600), or functionally-graded gyroid structures (FG600) were fabricated as illustrated in Figure 1. FG600 implants were created with a radially-varying pore size using a linear distribution of unit cell size to create a pore size distribution between 600 μm at the scaffold exterior and 300 μm at the scaffold midpoint. The porous fraction of all implants was designed with a height of 6 mm and Boolean merged to a slotted crown with a height of 1 mm prior to PBF, with a total implant diameter of 6 mm. The relative porosity of the G300 scaffold was estimated at 50% while the net porosity of the FG600 was estimated at 60%.

**Figure 1:**
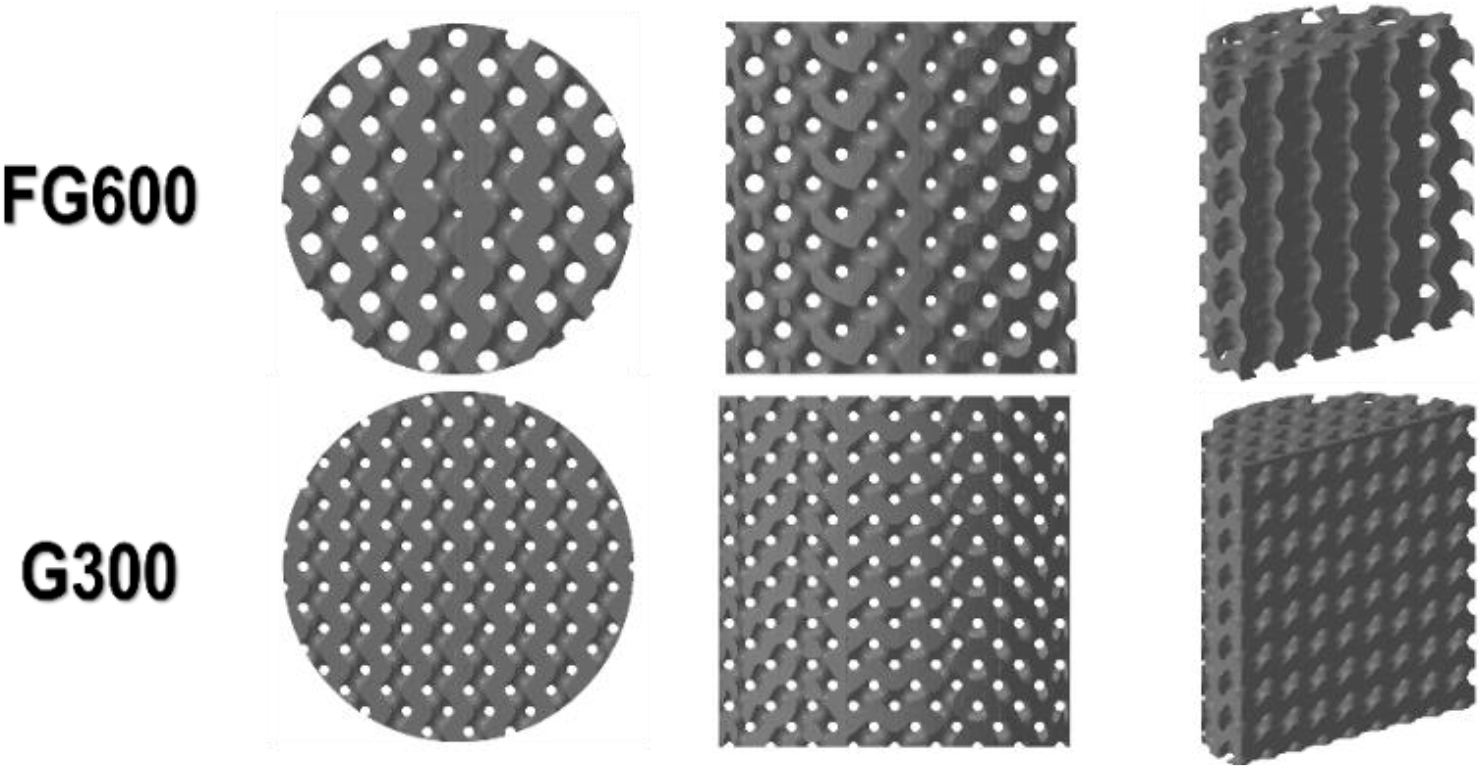
Overview of porous regions within the gyroid implants. The FG600 implant contained pores that decreased in size from 600 μm to 300 μm as a function of implant radius in the gyroid. The G300 implant contained a uniform array of 300 μm pores through the gyroid. The G600 implant (not shown) contained a uniform array of 600 μm pores through the gyroid.

### In Vitro Saos-2 Culture

Saos-2 cells (ATCC HTB-85) were seeded on uniform titanium gyroids to mimic osteoblast proliferation *in vitro* as described elsewhere [30]. Briefly, cells were seeded onto six replicates of G300 and G600 scaffolds using a seeding density of 10,000 cells/cm^2^ in a 24-well plate (approximately 17,700 cells per specimen). Culture was conducted in McCoy’s modified 5A media (Life Technologies Inc.) using incubation at 37°C and 5% CO_2_. After one, three, and seven days of culture, cell metabolism was measured by incubating samples with a mixture of 5% alamarBlue dye (Life Technologies Inc.) in culture media for 60 min and extracting the dye for fluorescence readings at excitation-emission wavelengths of 540/580 nm. A two-way analysis of variance (ANOVA) was conducted to determine statistical significance for each test group using R 3.6.1 *(α = 0.05)*. Following metabolic measurement at seven days, cells on a gyroid specimen were fixed by immersion in 0.25% glutaraldehyde in cacodylate buffer for 2 hr and stained with osmium tetroxide. Samples were dehydrated in a graded ethanol series (50%, 70%, 70%, 95%, 95%, 100%, 100%) in approximately 1 hr intervals before critical point drying in ethanol. Scanning electron microscopy (SEM) was conducted on the gyroid specimen following the application of a 5 nm platinum coating using a TESCAN VP microscope at an accelerating voltage of 10 kV to examine cellular interaction with the titanium.

### Surgical Implantation

20 adult male New Zealand white rabbits, weighing about 3 kg, were used in this study under the approval of the Animal Care Committee of Sun Yat-sen University (Approval No. SYSU-IACUC-2020-000198). Under general anesthesia and routine disinfection, a FG600 and a G300 implant were alternately and bilaterally placed in the left and right anteromedial, proximal tibial epiphyses of each rabbit to avoid bias from strong and weak responders [31]. The surface of the implant was set level with the cortical bone surface before suturing the surgical sites. Antibiotics were administered for three consecutive days after surgery. In order to trace and quantify new bone growth, calcein (C0875 Sigma-Aldrich) and alizarin red (A3882 Sigma-Aldrich) fluorochrome labels were injected into all rabbits in the 2^nd^ and 6^th^ week of implantation respectively. The doses of calcein and alizarin red were 5 mg/kg and 30 mg/kg respectively.

At each end-point (4 and 12 weeks after surgery), 10 rabbits were sacrificed, and the tibia-implant blocks were retrieved and fixed in 10% neutral buffered formalin solution. Five of the blocks were taken for X-ray microcomputed tomography (micro-CT) and the remaining five were used for the histological evaluation at each time point.

#### X-ray Microcomputed Tomography

Following tissue fixation, tibia-implant samples were subjected to an ethanol dehydration series (70%, 80%, 90%, 95%, 95%, 100%, 100%) in 48 hr intervals and gradually infiltrated with Embed812 resin (Electron Microscopy Sciences) in sequential ratios of 1:3, 1:1, and 3:1 with either acetone or propylene oxide. Resin was cured for 48 hr at 65°C. Micro-CT (Bruker Skyscan 1172) was conducted using a 100 kV beam and 100 μA current with a 0.5 mm Al filter. Voxel size during acquisition was set to 3.55 μm with a 0.25° rotation step. NRecon software (Bruker) was used to create an image stack from the projections using misalignment compensation, moderate ring-artifact reduction, and beam-hardening correction of 67%.

Image processing was conducted in Dragonfly 2022.1 (Object Research Systems) where all micro-CT datasets were subjected to a Gaussian smoothing operation with a kernel size of 9 and σ = 3.0. A 5-layered convolutional neural network (CNN) was created to semantically segment four classes within the micro-CT datasets (Figure S1), designated as implant material, bone residing within the implant, bone residing outside the implant, and resin/void space. The CNN was trained using augmented training data from 19 periodically sampled images across two separate datasets, where 80% of the training data was used for training and 20% of the training data was used for validation. Training was conducted using a patch size of 64, stride ratio of 0.5, batch size of 32, learning rate of 0.30, and was stopped early after 10 consecutive epochs without decrease in the validation loss to avoid overfitting.

Scaffold pore size validation and strut size variation was found by fitting a cylindrical sub-volume to the porous interior of the scaffold and masking regions attributed to implant and non-implant. Island removal was conducted to remove connected components with a voxel count of 1,000,000 or fewer and each region was assigned to a thickness mesh using a downsampling factor of three for three-dimensional measurement of pore and strut size. Bone volume fraction (BV/TV), measured here as the percent of available pore space in the implant filled with tissue, was measured by applying a cylindrical mask to the segmentation at the boundaries of the porous midsection and calculating the number of bone-classified voxels divided by the sum of total voxels (pore and bone). Percentage of bone-implant contact (BIC) was measured by performing a 3D cubic dilation with a kernel size of 15 on the region of interest (ROI) labelled as implant within the previously defined cylinder to create a zone within 25 μm of implant struts. The Boolean difference between the dilated ROI and original implant ROI was taken. BIC was then calculated as the ratio of the number of bone-labelled voxels within the Boolean subtraction divided by the total volume of the Boolean subtraction. BIC and BV/TV measurements were compared using a two-way ANOVA and Tukey’s HSD test using a significance value of α = 0.05.

### Scanning Electron Microscopy

Select bone-implant blocks were subjected to serial dehydration in acetone using 48 hr intervals and infiltrated with Embed812 resin in ratios of 1:3, 1:1, and 3:1 before transfer to 100% resin for curing at 60°C. Bone-implant samples were bisected using a low-speed diamond saw (Bueler Isomet) for backscattered (BSE) scanning electron microscopy. Specimens were polished (400-1200 SiC paper), mounted on an aluminum stub, and sputtered with 10 nm of carbon. Backscattered electron images were acquired using a JEOL 6610 LV microscope (JEOL, Peabody, USA) at an acceleration voltage of 10 kV.

To reveal the underlying osteocyte network of samples, approximately 5 ml of 37% phosphoric acid was placed on the top surface of the embedded samples for 10 seconds. The samples were then rinsed with excess deionized water for 20 seconds, immersed in a bath of 5% sodium hypochlorite for 5 minutes, thoroughly rinsed again for 30 seconds, and air-dried overnight. The samples were prepared for SEM imaging with a 10 nm gold coating and imaged using secondary electron imaging in a JEOL 6610LV microscope (JEOL, Peabody, USA) at 7 kV.

### Histology

Tibia-implant specimens for histology were dehydrated using increasing concentrations of alcohol (70%, 80%, 90%, 95%, 95%, 100%, 100%) and then embedded in methylmethacrylate for sectioning. Cutting of 100-μm-thick sections, parallel to the long axis of the implants, was performed by using an EXAKT E300CP (EXAKT, Germany) to extract a single slide section from the midpoint of the implant. The sections were then ground by EXAKT 400CS (EXAKT, Germany) and stained with hematoxylin and eosin (H&E).

H&E-stained histological sections were observed using an EVOS inverted microscope (Life Technologies Inc.) to view cell and bone matrix deposition at ROIs defined on the interior and exterior of the scaffolds. Four slides for each implant type at 4 wk and 12 wk of implantation were imaged using bright-field imaging, and fluorescent signals from calcein and alizarin red on the same sections were observed using an A1R HD25 inverted confocal microscope (Nikon) to assess locations of calcification in the new tissue after 2 weeks and 6 weeks. Fluorescent images were overlaid in Dragonfly 2022.1 with gamma correction and implant boundaries were found using a grayscale-derived segmentation of the implant volume.

## Results and Discussion

### In Vitro Scaffold Characterization

*In vitro* characterization of the gyroid scaffolds (G300 and G600) was conducted using the Saos-2 cell line as a rapidly proliferating approximation of the behaviour of osteoblasts on the scaffolds [32]. By measuring the metabolic activity of Saos-2 cells on the scaffolds at one, three, and seven days of culture (Figure 2A), statistically significant growth was observed between each time point regardless of scaffold geometry. At each unique time point in the seven-day study, gyroid scaffolds in the G300 and G600 groups showed similar potential for osteoblast proliferation and *in vitro* bone growth via metabolic measurement. SEM of cellular interactions with the additively manufactured titanium showed that Saos-2 cells adhere to the Ti6Al4V substrates and can co-locate to bridge gaps between partially-fused titanium powder particles on the surface (Figure 2B).

**Figure 2:**
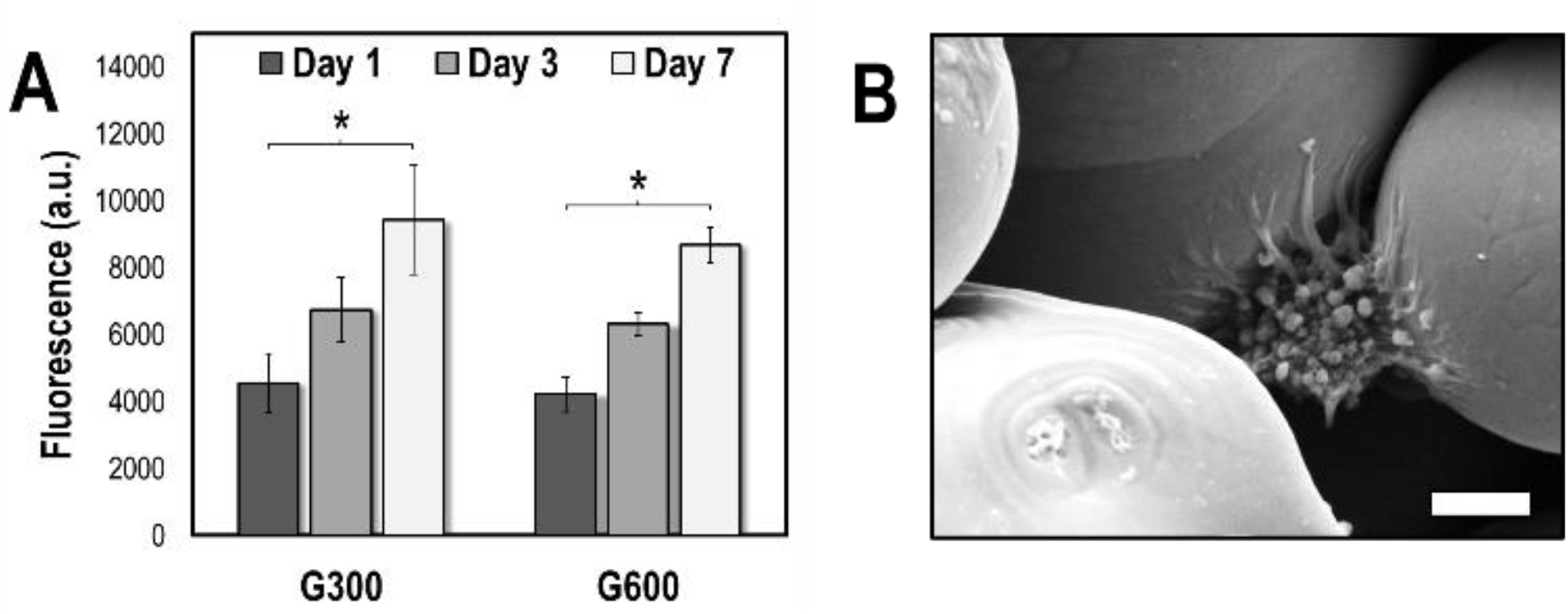
In vitro assessment of uniform gyroid Ti6Al4V scaffolds. (A) Saos-2 metabolic activity increases for each type of scaffold at three and seven days of culture (p < 0.05). Scaffolds showed comparable behaviour with pore sizes of 300 μm and 600 μm at each time point. (B) Adherent Saos-2 cell on the surface of an additively manufactured Ti6Al4V gyroid specimen, showing interaction with the cleft between two sintered powder particles on the surface. Scale bar: 5 μm.

For laser powder-bed fusion additive manufacturing, in the absence of post-processing surface modification strategies such as acid-etching [33,34] or grit-blasting [35,36], the packing behaviour of surface-bound metallic particles is governed by characteristics of the feedstock powder and processing parameters defined during selective laser melting [37]. Where similar observations of *in vitro* cell-surface interaction have been observed on stainless steel powder particles [38], it seems as though the pore design of the scaffold may have less of a role than physical or chemical modification of the surface for *in vitro* proliferative effects. Similar fluorescence readings were obtained for the G300 scaffolds and G600 scaffolds, despite the increase in internal surface area in the G300 group. It appears the bulk of the early cellular activity can therefore be attributed to near-surface cells in the scaffold. Similar behaviour has been observed with 3D cell culture in porous microcarriers for other cell types, including chondrocytes [39] and adipose-derived stem cells [40], where cell proliferation to internal surfaces only starts to become evident around five or seven days of culture.

### ray Microcomputed Tomography

#### Scaffold Feature Size Validation

To assess the deviation in pore and strut size in each *in vivo* scaffold type (G300 and FG600) from the design, measurements were extracted from micro-CT scans of the implants (Figure 3).Representative slices from the porous midsection (Figure 3A-B) show a distinguishable difference between the two scaffolds (3A: G300 uniform, and 3B: FG600 functionally-graded) at a glance based on their relative pore size. Mapping the pore and strut size distributions for the G300 implant (Figure 3C and 3E) showed a similar distribution in each. The highest frequency of pore size for the G300 implant was measured to be between 300-310 μm, indicating that the build parameters utilized during additive manufacturing result in parts with a relatively high tolerance. The median pore size for the G300 implants lies in the range of 250-260 μm, however, the lower bound of the distribution is subject to image processing artifacts as pores at the edge of the scaffold can be partially truncated when performing the Boolean intersection with the bounding cylinder or some marginal pore occlusion by partially-fused powder particles. Similarly, the most frequent strut size in the G300 scaffold also ranges between 300-310 μm with a median between 250-260 μm. For uniform pore sizing, the gyroid topology appears to offer a consistent interconnected pore structure throughout the scaffold interior and avoids sharp self-intersections as is typical of TPMS geometries.

**Figure 3:**
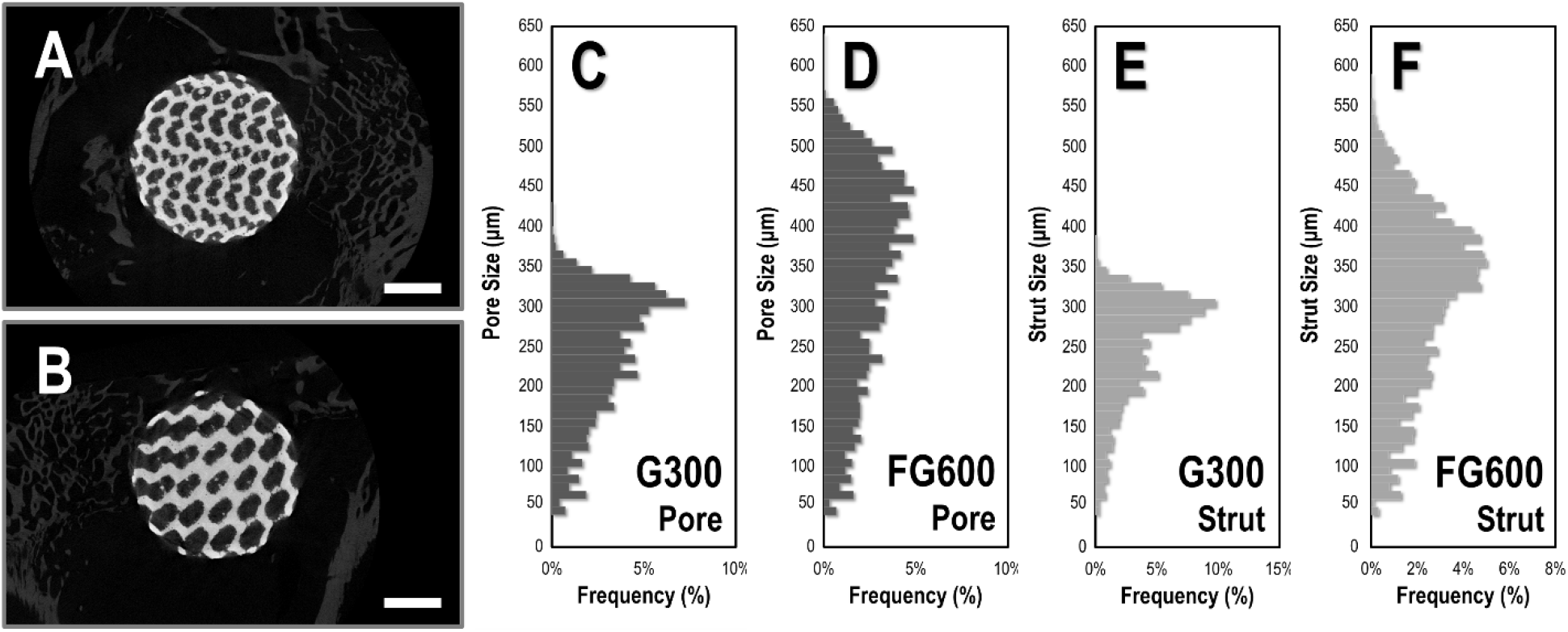
Micro-CT and gyroid topology validation. (A) Micro-CT slice showing cross-section of G300 scaffold after 12 wk of implantation in leporine tibia. (B) Micro-CT slice showing cross-section of FG600 scaffold after 12 wk of implantation in leporine tibia. (C) Pore size distribution (n = 10) within the G300 scaffold interior. (D) Pore size distribution (n = 10) within the FG600 scaffold interior. (E) Titanium strut size distribution (n = 10) in the G300 scaffolds. (F) Titanium strut size distribution (n = 10) in the FG600 scaffolds. FG600 scaffolds appear to have a wider distribution of both pore diameter and strut diameter as a result of the functionally-graded pore design. Scale bars: (A-B) 2 mm.

In contrast, it is clear from the pore and strut size distributions in the FG600 implant that the addition of functional grading widens the overall pore and strut size distribution in the implant. Here, larger pore sizes are more commonplace, with the highest frequency between 440-450 μm although the median lies between 350-360 μm. Due to the relative size of the implants, the functional grading from 600 μm to 300 μm occurs over a short radius of 3 mm in the scaffold. As a result, only the outermost surface of the scaffold is anticipated to have pores of 600 μm diameter before shrinking with radius and this is the portion that is most prone to edge effects during image processing. The titanium strut diameter has a slight shift towards smaller values in the FG600 scaffold, with a mode between 350-360 μm. The mismatch between pore size and strut size distributions in the FG600 implant can be attributed to the overall scaffold design. Since functional grading of the scaffold is with respect to the entire unit cell, it is possible to tune topology without one-to-one inverse proportionality in the strut-pore relationship.

#### Bone Apposition

Bone apposition occurred within the pores and surrounding the exterior of both scaffolds at 4 and 12 weeks in the tibia, as demonstrated in the micro-CT reconstructions in Figure 4. The BV/TV for the G300 implants at 4 and 12 wk was measured at 0.68 ± 0.10 and 0.74 ± 0.04, respectively. For the FG600 implants, BV/TV was measured at 0.58 ± 0.12 after 4 weeks and 0.47 ± 0.04 after 12 weeks. As illustrated in Figure 5, a significant difference *(p < 0.05)* was found between the two types of scaffolds after retrieval, with the functionally-graded scaffold (FG600) containing a lower bone volume fraction than the scaffold with uniformly-sized pores (G300).

**Figure 4:**
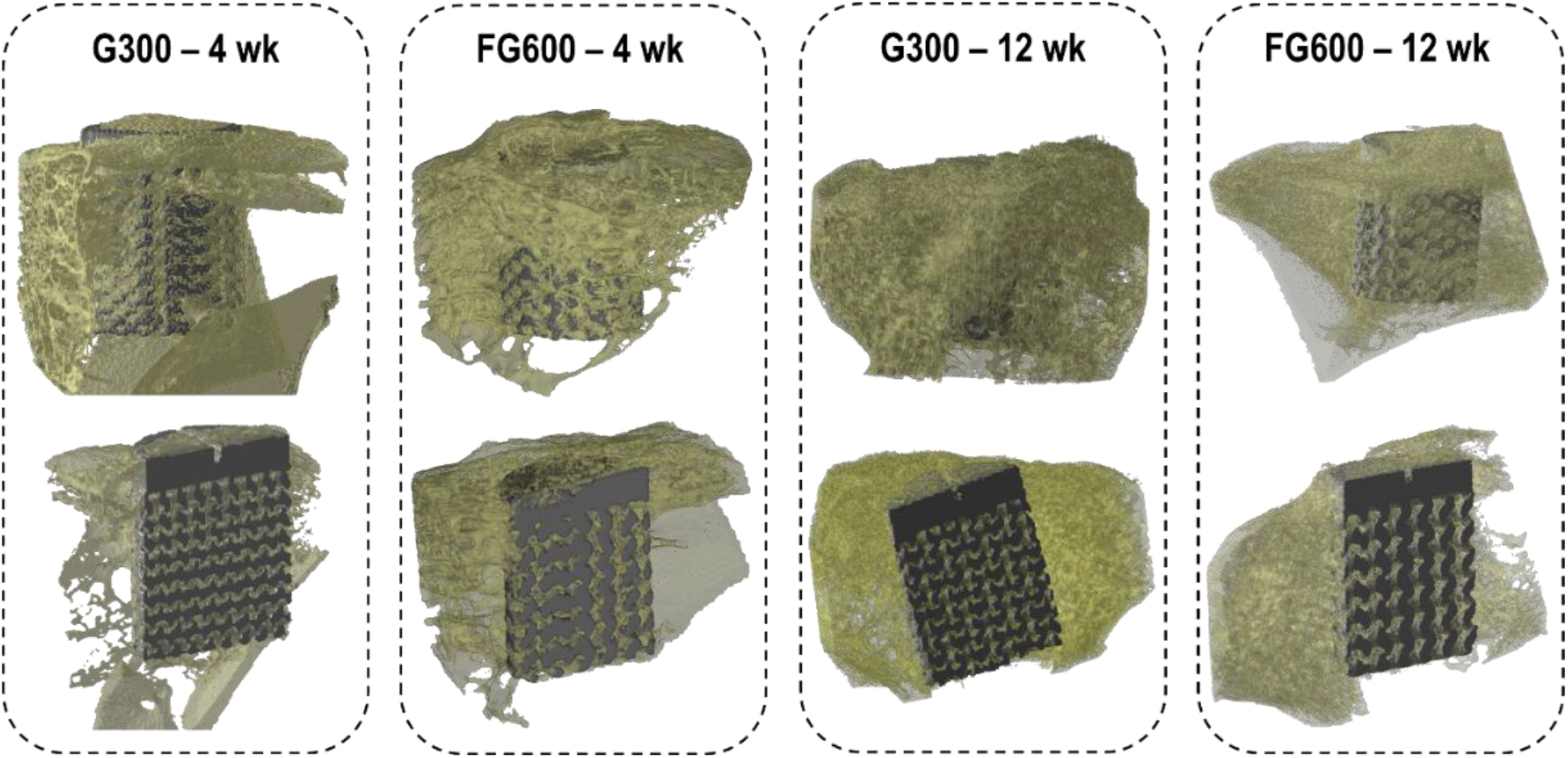
3D Micro-CT reconstructions at 4 wk and 12 wk. Formation of bone inside the tibial defect area has occurred after four weeks and twelve weeks, resulting in implant fixation. The porous region of the implants is situated within the trabecular fraction of the epiphysis in the leporine tibia, while the full-density titanium in the crown lies in the cortex. Longitudinal cross-sections for all implants show evidence of bone apposition at both four weeks and twelve weeks.

**Figure 5:**
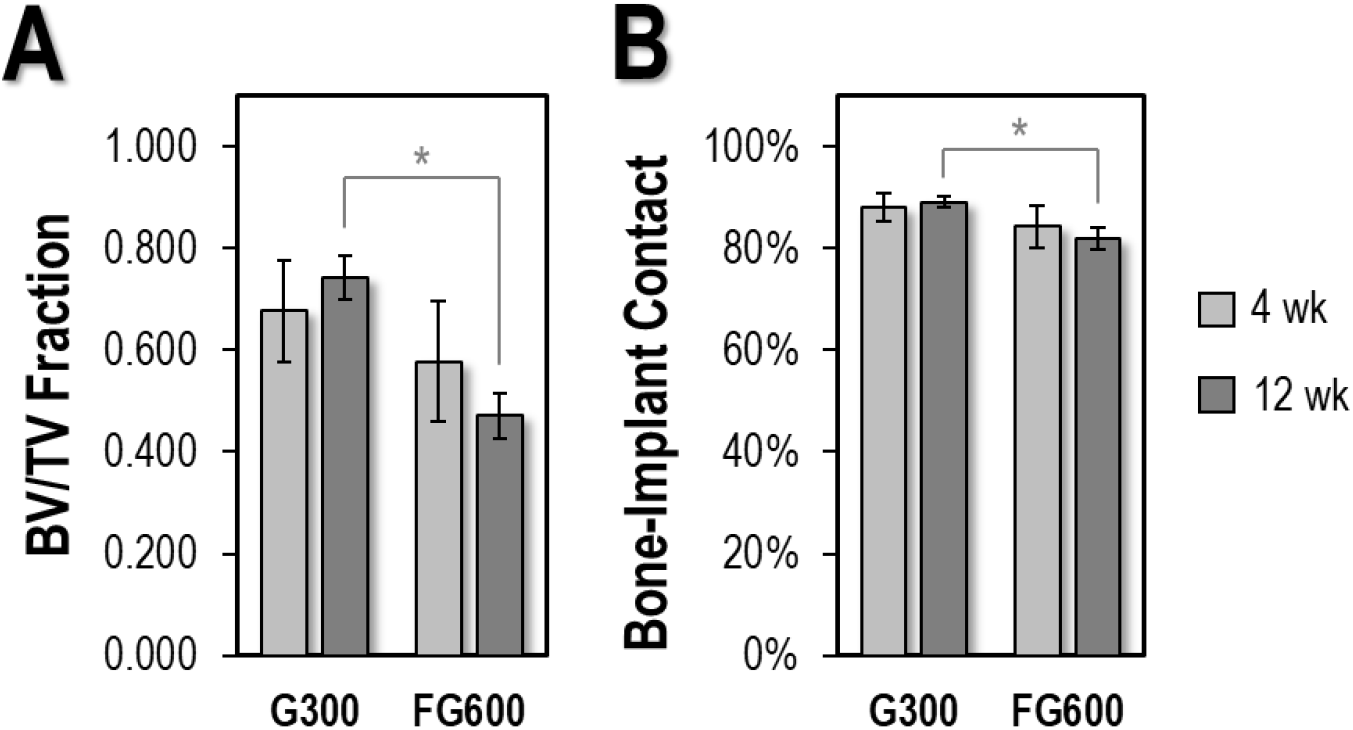
Histomorphometry of G300 and FG600 scaffolds at 4 wk and 12 wk. (A) Bone volume fractions for each scaffold. The uniform 300 μm pore size retains a greater bone volume fraction at 12 wk than the implant with functionally-graded pore size. (B) Percentage of bone-implant contact for each scaffold. BIC is greater at 12 wk in the implant with a uniform 300 μm pore size.

The measured BIC values for the G300 implants at the 4-week time point were 87.9% ± 2.7%, with a similar value after 12 weeks (89.1% ± 1.1%). Ranges in the FG600 implants were approximately the same amount, with 84.2% ± 4.1% BIC at four weeks and 82.0% ± 2.2% at twelve weeks. The G300 implant had significantly greater BIC at the twelve-week endpoint compared to the FG600 implant *(p < 0.05)*. BIC measurements in these porous scaffolds appear greater than what has been reported for pore sizes of 800 and 1200 μm after 3 and 8 weeks in leporine femora using scanning electron microscopy (SEM) [41], but it is also important to note that differences between the studies could also be associated with differences in resolution and measurements by 2D SEM and 3D micro-CT. Micro-CT of porous titanium with a pore size range of 500-700 μm placed into ovine femora yield BIC measurements of approximately 60% at 26 weeks [42]. Imaging artifacts can occur at the implant interface due to attenuation from the titanium [43]; using a threshold of 25 μm from the implant surface for BIC assessment may help reduce noise in these interfacial calculations, but the number of transitions from bone to implant along the path of a single X-ray is still expected to be greater in implants with more pores in the cross-section. Correlation to histological observation is important to confirm trends observed through micro-CT at the bone-implant interface.

Backscatter electron imaging in the SEM was also performed on cross-sections of the implant to assess the degree to which bone interacts with both types of implant. Based on a perspective from a lower magnification (Figure 6A-B), bone formation is most apparent at the crown of the implant and among the outermost implant struts. Examination by SEM into the interior of the implants suggests an overestimation of BIC and BV/TV measurements in the micro-CT due to beam hardening, but close bonding between bone and implant is still visible towards exterior scaffold sites at both 4 and 12 weeks (Figure 6C-D). Resin cast etching reveals underlying osteocytes and cell processes in the newly formed peri-implant bone (Figure 6E-F). Roughly 20 μm from the interface of a FG600 implant, the osteocytes are co-aligned with their major axis parallel to the titanium surface. An intervening layer of mineralized matrix separates the osteocytes from the titanium. As secretion of extracellular matrix from osteoblasts is a polar phenomenon, it is likely that these osteocytes were once adherent to the outermost surface of the implant and deposited osteoid as a template for early bone formation at the scaffold exterior. Osteocytes further from the implant have a rounded appearance, and potentially a higher surface area to volume ratio. Among the near-implant osteocytes (Figure 6F), cell processes appear majorly oriented into circumferential arrays with connecting branches oriented tangentially.

**Figure 6:**
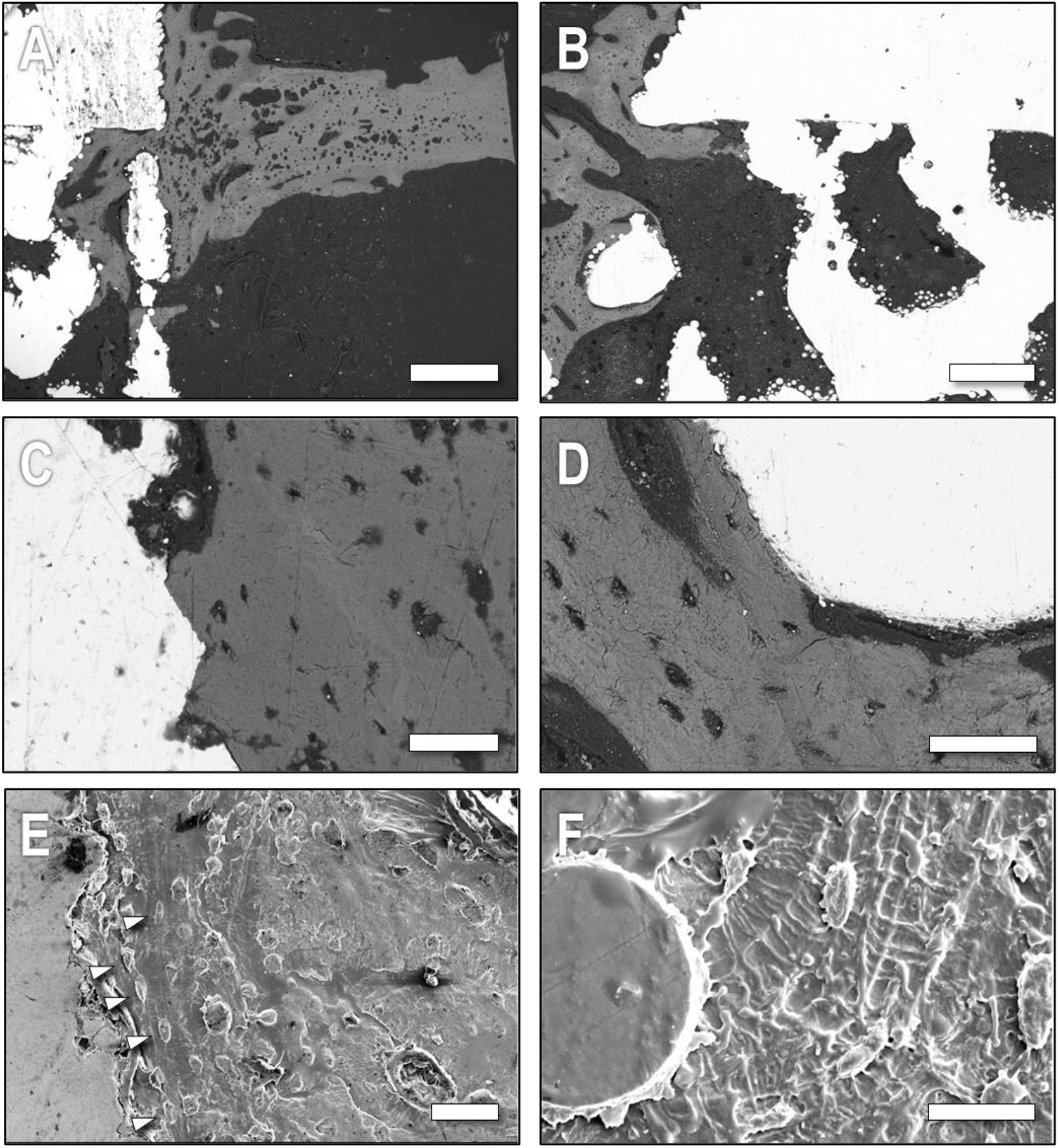
Scanning electron microscopy of the bone-implant interface. Bone formation at the exterior of (A) G300 and (B) FG600 scaffolds after 12 wk of implantation. Higher magnification view showing close bone-implant contact after both (C) 4 wk in a G300 scaffold and (D) 12 wk in a FG600 scaffold. (E-F) Following resin cast etching, osteocytes at the interface of a G300 implant after 12 wk have an oblong morphology (arrowheads) but become rounded with increasing distance. Propagation of the lacunocanalicular network appears in both radial and tangential patterns around the near-implant osteocytes. Scale bars: (A-B) 500 μm; (C-E) 50 μm; (E) 20 μm.

### Histology of Scaffolds in Leporine Tibiae

#### H&E Staining

Representative histological sections detailing interior and exterior sites of FG600 and G300 implants retrieved at 4 wk are shown in Figure 7. Additional H&E sections can be found in Figure S2 and S3.

**Figure 7:**
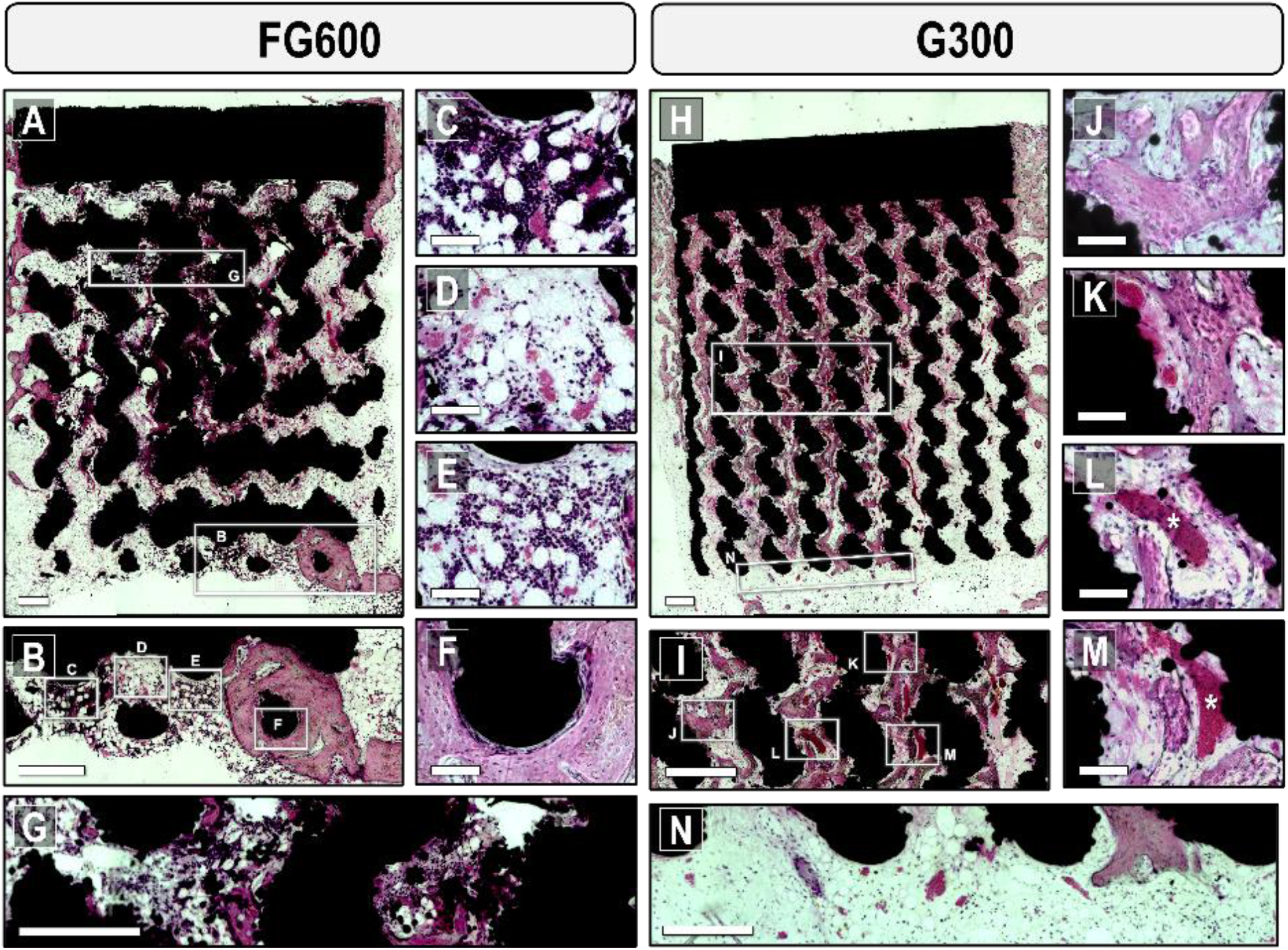
Histology of interior and exterior implant sites after four weeks for an FG600 (A-G) and G300 (H-N) implant. Hallmarks of bone formation throughout the FG600 scaffold with H&E staining after four weeks of implantation. (B) Inflammatory interactions occur at the FG600 exterior with a trabecular and marrow-like appearance around the initial clot, including the formation of: (C) Dark-appearing cells, likely lymphocytes, apparent among clusters of adipocytes and erythrocytes; (D) Erythrocyte-rich regions in the granulation tissue with fewer white blood cells present; (E) Inflammatory tissue without formation of a mineralized matrix at the implant interface and an intermediate fraction of white blood cells; (F) More extensive bone regeneration to create intimate bone-implant contact at the scaffold exterior. (G) Formation of bone matrix along with inflammatory tissue occurs within the FG600 interior. (H) Patterns in early osseointegration of the G300 implant. (I) Thin regions of appositional bone tend to span the implant interior with less inflammation than in the FG600 implants. (J) Bridging of the scaffold struts occurs with the formation of new trabecular bone. Bone appears to integrate closely at the strut surface and partially contour around titanium particles. (K) Competition occurs at the strut surface between microvascular entities and osteocyte-rich bone. The orientation of new bone seems to be mostly in parallel with the pore wall. (L-M) Vascular structures, denoted by asterisks, are able to permeate deep into the scaffold interior with bone forming around the vasculature. Microvasculature propagates through both the void volume in the scaffold and along the titanium surface. (N) G300 exterior with high content of adipocytes, a network of microvasculature, and low relative counts of white blood cells. Scale bars (A-B, G-I, N) 500 μm; (C-F, J-M) 100 μm.

At the outermost layer of the FG600 scaffold (Fig. 7B), there is evidence of an inflammatory response occurring in close proximity to the apex of the titanium scaffold. The inflammatory tissue that has formed varies widely in its composition across distances of 500 μm or less. In the inset shown in Figure 7C, dense clusters of dark-coloured basophilic cells or white blood cells tend to aggregate around dispersed lipids in the tissue. Due to their lack of eosinophilic cytoplasm in most cases and large nuclear fraction, these cells are consistent with lymphocytes [44]. At the FG600 apex, the white blood cells (black/purple) and erythrocyte-rich microvasculature (red) both infiltrate the space between lipids (white) in the region. This region has direct line-of-sight access to the scaffold exterior, with no titanium struts obfuscating osteoblasts or other biological response. In the first interior layer of pores (Figure 7D), there is a much lower density of white blood cells present in the inflammatory tissue, with similar volumes of red blood cells in the titanium-bound microcapillaries and a comparable dispersion of lipids near the titanium surface. The immune cells here appear to have a higher cytoplasmic fraction than in the high-density region and are more consistent with granulocytes [44]. Figure 7E shows a neighbouring region, with a finer capillary structure in the tissue and a slightly higher granulocyte density compared to what is seen in Fig. 7D. With respect to the apposition of bone tissue at the implant surface, nearby regions have an ‘interlocking’ effect where new bone has formed to fully encapsulate an exterior scaffold strut (Figure 7F). Osteocyte lacunae are visible within the bone matrix as close as 20 μm from the implant surface, with an approximate concentric alignment around the contours of the strut surface. The extent of bone regeneration at this site and maturity of the tissue seems to indicate that bone at exterior sites is further along in the modelling cascade than bone nucleating at interior sites. This distinction is often made in porous implant materials, where quantitative histomorphometry can be subcategorized into interior and exterior moieties depending on whether near-surface pores are sampled [42].

While the *in vitro* results at early timepoints showed a biological response that was predominantly occurring on the exterior of the scaffolds or within the first layer of pores, the *in vivo* results showed formation of bone in the scaffold interior after only four weeks of implantation. Figure 7G and 7I illustrate bone regeneration at interior sites within the scaffolds at this time point, where lower-magnification images show a disperse network of thin bone that populated the scaffold interior in the G300 implant (Figure 7H-I) and residual inflammatory tissue mixed with bone matrix in the FG600 implant (Figure 7I). The apical portion of the G300 implant (Figure 7N) after four weeks had a characteristic appearance of bone marrow and is mostly filled with adipocytes and microcapillaries rather than white blood cells. At the crown of the G300 and FG600 implants, osteoconduction from cortical regrowth was beginning to occur, providing inward growth into the pores nearest the cortex.

Looking at pores at increasing radial depth into the G300 scaffold (Figure 7I) tends to illustrate a lack of depth-dependency for bone growth, with pores at the midpoint of the G300 scaffold containing similar amounts of bone to shallow pores near the implant surface in this instance. The location of the scaffold in the medullary canal also appeared to affect bone tissue apposition, as osteogenesis appeared more dominant in the leftmost and central regions of Figure 7G for the G300 implant. However, at sites within the scaffold interior, appositional bone tissue appeared to bridge adjacent scaffold struts at the four-week endpoint with an appearance that is consistent with trabecular bone (Figure 7J-K). Osteocytes in these regions of new bone formation appeared mostly disorganized, with high density and no clear direction of co-alignment as is typically seen in mature bone tissue [45], e.g., the apical regrowth in the FG600 implant. Tissue regeneration in the scaffold was not always unidirectional with respect to the scaffold struts, where bone formation occurred in a predominantly transverse direction (as in Figure 7J), or with longitudinal components (Figure 7K). There were also fewer lipids and granulocytes in and around the G300 scaffold compared to the FG600 implant in Figure 7, suggesting that bone repair and the fracture healing cascade may be slightly more advanced in the G300 implant compared to the FG600 implant.

In terms of biological interaction, a fine network of vasculature permeated throughout the entirety of the implant to aid in both tissue formation and homeostasis. These microcapillaries contained an abundance of erythrocytes, denoted by asterisks in Figure 7L and 7M, and appear to be sectioned obliquely based on their aspect ratio. The ability of this provisional capillary network to maintain carrying capacity is important for osseointegration as it mediates the delivery of oxygen, growth factors, and the removal of waste by-products at these interior scaffold sites during bone modelling and remodelling [46]. With respect to osseointegration, neovascularization has been documented to occur on the surface of solid titanium implants long before the formation of new bone tissue [47] with local expression of factors such as Runt-related transcription factor 2 [48] that control the pre-osteoblast to osteoblast transition. Here, we note that this newly formed vasculature was able to infiltrate a porous metallic scaffold in leporine tibial defects in a similar manner to what has been seen for murine cranial defect interfaces with solid titanium [47]. The newly formed vasculature has a tendency to interface and propagate along the porous titanium (as in Figure 7K and 7M) but also spans scaffold struts in a freeform manner (as in Figure 7L). At early timepoints, the capillary network is observed to periodically interact with the titanium as a self-supporting mechanism during trabecular modelling.

Examination of the FG600 (Figure 8A-F) and G300 (Figure 8G-L) scaffold geometries after twelve weeks shows evidence of sustained osteogenesis between the four and twelve-week endpoints. The interior pores of scaffolds retrieved after 12 wk tended to have much higher adiposity than those retrieved at 4 wk, where the bulk of the tissue inside the scaffold was consistent with that of myeloid tissue (Figure 8B and 8L) with radially-oriented eosinophilic regions characteristic of the bone matrix that bridge neighbouring struts in the gyroid topology. The consistent patterns in bone apposition within the scaffold interior mirrored what was seen in the micro-CT reconstructions, confirming that the unit cell topology can, at least in part, govern the directionality of nascent tissue in the scaffold interior. At 12 wk, adaptation of the tissue is noted to maintain this pattern and preserve the functional connection of bone and implant throughout the scaffold interior. It is possible that the increase in adipocyte-rich myeloid tissue within the scaffold (Fig. 8C-E) can be attributed to a shift in stem cell differentiation, where osteoblast progenitors can ostensibly commit to an adipogenic pathway [49,50]. As bone marrow is formed inside the scaffold interior, hematopoiesis can occur to sustain red and white blood cell production. Figure S4 shows an interior pore channel, with a myeloid structure, including the presence of a suspected megakaryocyte along with white blood cells, red blood cells, connective tissue, and adipocytes – indicating that residual platelet production and clotting [51] was still occurring in the scaffold after 12 wk. Comparing the two types of scaffolds, bone matrix appeared more densely distributed throughout the interior of the G300 implant due to the smaller pore size helping to facilitate bone formation that spanned multiple scaffold struts.

**Figure 8:**
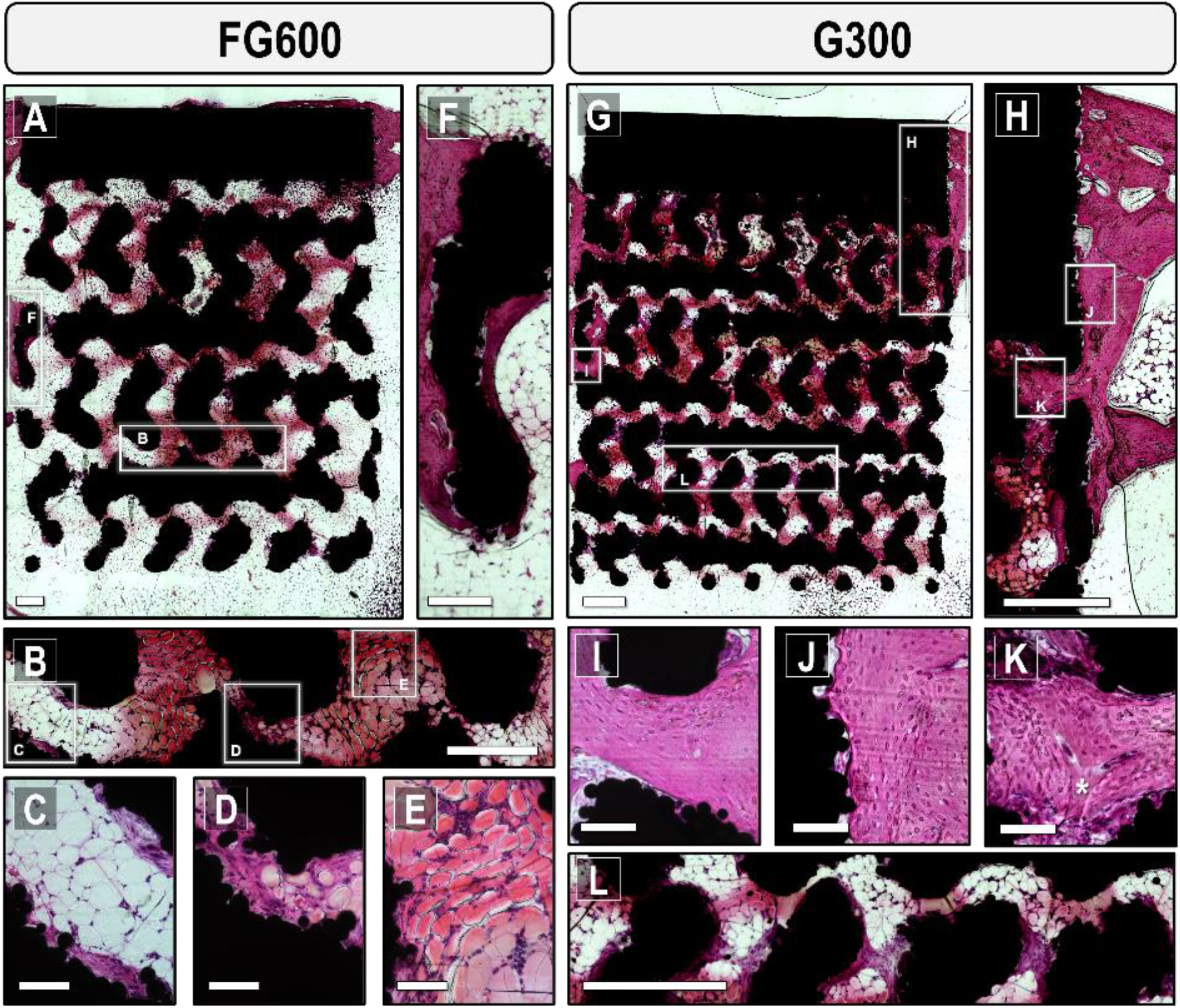
Histology of interior and exterior implant sites after twelve weeks. (A) Bone formation throughout the FG600 scaffold with H&E staining after twelve weeks of implantation. Appositional bone has coarsened compared to the four-week endpoint and fully occupies the pore diameter in some instances. (B) Inset in the FG600 interior shows high lipid content, with: (C) Circular adipocytes in the centre of the pore, flanked by regions of bone matrix along the titanium; (D) Predominantly bone matrix where pores narrow; (E) Elliptically-shaped adipocytes in an eosinophilic matrix, with marginally higher white blood cell clustering compared to other myeloid regions. (F) Bone integration at the scaffold exterior, with bone fully encapsulating an outer strut. (G) Bone formation throughout the G300 scaffold with H&E staining after twelve weeks of implantation, with a seemingly higher bone area fraction than the FG600. (H) Inset at the crown of the implant shows cortical osteoconduction beyond to the first layer of pores within the scaffold. (I) Newly formed tissue in the trabecular region enters the implant, with mature osteocytes that concentrically align with the upper scaffold strut. (J) The tissue formation front at the crown tends to only interact with a single tangential region on each sintered powder particle. (K) New bone tissue completely surrounds a branching microcapillary, denoted by an asterisk, where the capillary is roughly located at the midplane of the two scaffold struts. Scale bars (A-B, G-H, L) 500 μm; (F) 250 μm; (C-E, I-K) 100 μm.

There was generally less inflammatory tissue present near the implant exterior at twelve weeks for both scaffold types, although some persisted near the apex of the implant and there were fewer white blood cells present. Similar to trabecular integration at four weeks, there were regions (Figure 8F) where mature bone wrapped around the entirety of the outermost scaffold struts. At twelve weeks, newly formed bone in the trabecular and cortical regions tended to be coarser. Bone within the trabecular region was able to branch into the porous interior, spanning the entire diameter of the pore in some instances. Osteocytes within these inwardly-directed trabeculae had a mature phenotype according to their shape [52], with a flat lacunar morphology (Figure 8I). These osteocytes tended to align more closely with the upper scaffold strut in this pore, perhaps due to the higher number of sintered powder particles and therefore rougher nature of the lower strut. Tissue integration in this trabecular region followed the contours of surface-bound titanium particles but this was not ubiquitous on the implant surface. Cortical regeneration along the implant crown showed that the bone formation front can instead form tangential to the surface particles, with no tissue penetrating the crevices between powder particles (Figure 8J). Osteocyte lacunae in this region of the crown were also consistent with a mature morphology, with their flatter edge parallel to the implant surface. In some instances, at four and twelve weeks, a clear path of osteoconduction can be observed where bone developed at the existing cortex and grew along the crown of the implant to merge with other regions of newly formed bone. This bone then breached the porous layers adjacent to the cortex to enter the scaffold interior – a phenomenon observed in both the FG600 and the G300 scaffolds.

After twelve weeks, microvascular capillaries no longer appeared to be interfacing with the titanium scaffold. Capillaries instead appeared with typical appearance in mature bone tissue, propagating through continuous vascular canals in the mineralized bone tissue, with one such example shown in Figure 8K. The vascular structure seemed to follow the gyroid scaffold topology, bifurcating at a site where the pore network branches. The vascular entities also appeared to be roughly centred at the midplane between neighbouring scaffold struts. Preservation of these capillaries throughout the scaffold is essential to allow for survival of bone at the innermost scaffold locations by acting as a transport system for biological macromolecules and regulating tissue metabolism.

It may indeed be worthwhile in the future to assess the spatial distribution of factors such as Runt-related transcription factor 2 (RUNX2) or peroxisome proliferator-activated receptor-γ (PPARγ) [53] to validate the decline in osteogenic commitment inside the scaffolds and corresponding increase in adipocyte formation. This would provide information about the differentiation of mesenchymal stem cells over time inside the pore network of a scaffold and provide a basis for developing temporally-relevant physical and chemical surface modification strategies.

#### Fluorochrome Labels

To compare sites of calcification throughout the interior of the implant, fluorochrome signals were compared using calcein injection at two weeks and alizarin red at six weeks post-implantation. By comparing fluorochrome-labelled regions in these implants with H&E-stained sections (sampled at four weeks and twelve weeks), it is possible to localize regions of matrix mineralization within the tissue. For all implants in Figure 9, a fluorescent signal from calcein was evident within the scaffold interior after only two weeks, indicating that a framework for mineral transport into the scaffold was present in a very early period following implantation. The co-localization of calcein signal at two weeks and tissue present at four weeks (Figure 9E-H) indicates that similar quantities of mineralized tissue were present in the scaffold interior at early timepoints, despite the more extensive network of bone matrix in the G300 implant. The same trends persisted for the implants retrieved at twelve weeks, where the bulk of the fluorescent signal was co-localized with stained regions in the H&E sections. These early mineral loci did not appear ubiquitously through the matrix material and did not yet span the full pore diameter in regions where they appeared, instead nucleating as smaller clusters.

**Figure 9:**
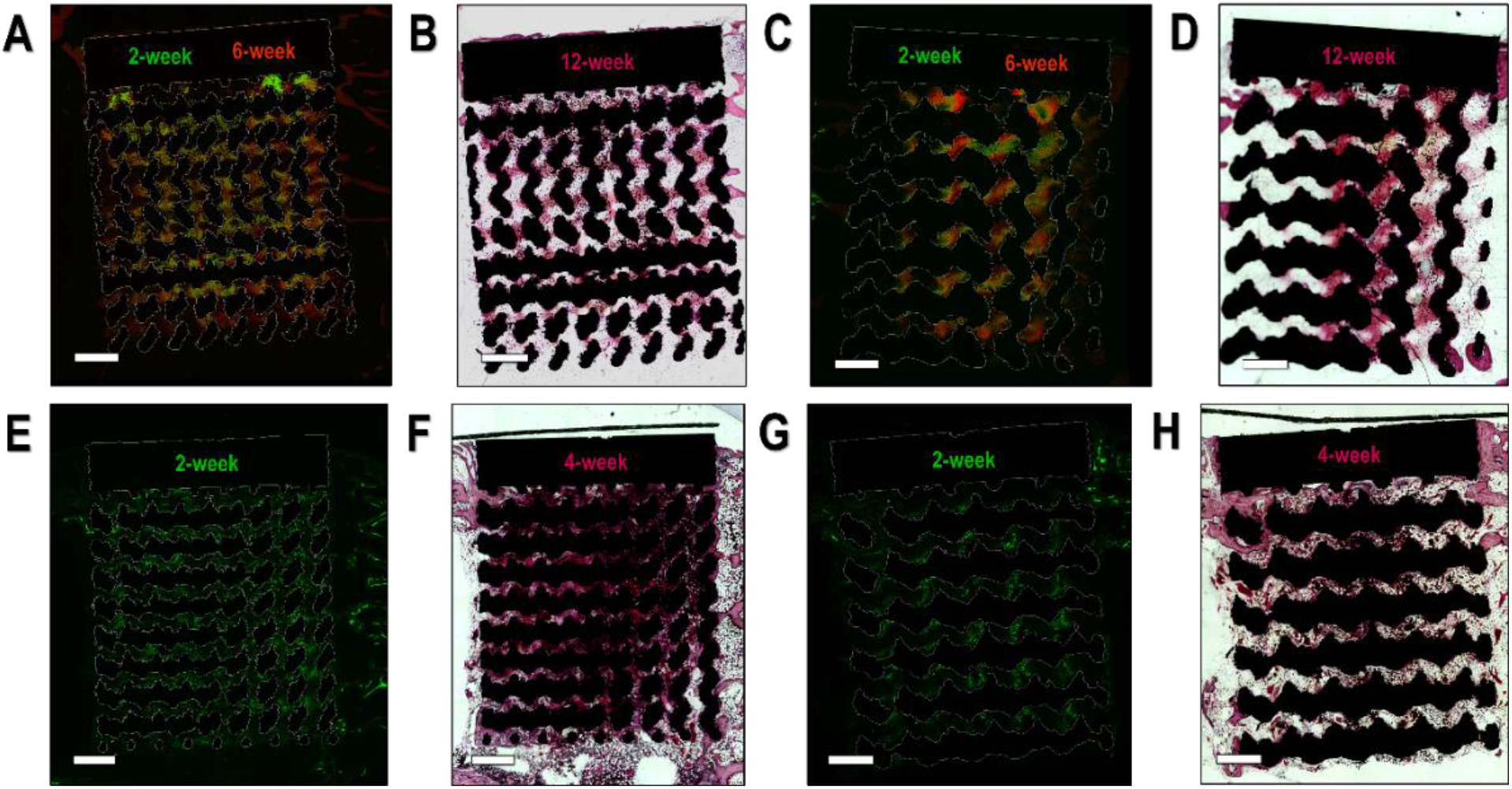
Fluorochrome labelling of calcification loci. (A-D) Fluorochrome images of calcein injected two weeks post-implantation, alizarin red injected six weeks post-implantation, and correlative H&E-stained section after twelve weeks in the (A-B) G300 and (C-D) FG600 implants. Calcified regions expand between the two-week and six-week endpoints, correlating closely with tissue in the H&E section at twelve weeks. (E-H) Calcein fluorochrome and H&E stain showing correlative bone apposition at two and four weeks in (E-F) G300 and (G-H) FG600 implants. Fluorescent signals are bright throughout the implant interior where bone matrix is present, indicating that mineral transport is consistently occurring within the scaffold interior within the first two weeks and corresponds with regions of connective tissue in the H&E micrographs. Scale bars 1 mm.

In both the H&E and fluorochrome images, periodicity was evident with respect to where new tissue formed. At similar radii through the implant, there were consistent patterns in appositional bone formation inside the implant. Bone that spans two neighbouring struts of titanium had a consistent and somewhat unidirectional appearance in all pores on the same azimuth (i.e. along the same ‘rows’ of pores in Figures 8 and 9). For the FG600 implants, this appositional bone had a tendency to orient itself in the radial direction of the implant, while the G300 implants tended to have a more oblique direction in the tissue. Pobloth et al. described a similar ability for bone to grow through axial, perpendicular, and oblique pore directions along the interior struts of titanium scaffolds, using collagen directionality to assess this form of tissue patterning [54]. Although the unidirectionality may be an artifact of sectioning orientation, commonalities in appositional bone orientation in the fluorochrome and H&E deserve further investigation. Where unit cell topology was shown to have only a minor role for *in vitro* scaffold assessment, the *in vivo* results suggest that topology can indeed play a larger role with respect to the directionality of osteogenesis in the scaffold interior.

The evolution of mineralization from two weeks to six weeks was more evident in the FG600 implants, where calcified regions appeared to propagate outward from the original loci present at two weeks (Figure 9C). This was consistent with the use of fluorochrome labelling in other studies, where early endpoints tended to have smaller fluorochrome-labelled areas near the interface of a titanium implant [55]. Alizarin red fluorescence showed that mineralized tissue can span across entire pores, effectively linking two titanium struts. The G300 group of implants showed a similar trend where existing mineralized tissue coarsens, but there were also regions where new mineral apposition seemed to occur at six weeks, connecting scaffold struts with no prior mineral adjacent to them.

## Conclusions

Additive manufacturing of titanium enables the production of complex scaffolds with an interconnected pore geometry to influence early osseointegration and short-term implant stability. Ti6Al4V gyroid structures displayed *in vitro* cytocompatibility with respect to Saos-2 metabolism. *In vivo* assessment of the uniform and functionally-graded gyroid structures in leporine tibiae both show potential for improving early implant outcomes by encouraging osteogenesis to occur on the scaffold exterior and interior. In general, newly formed bone spans internal scaffold struts with a common directionality relative to neighbouring regions of bone apposition, governed at least in part by the strut orientation of the scaffold itself. BV/TV and BIC measurements from micro-CT reconstructions demonstrated that values were greater for G300 implants compared to FG600 implants at 12 wk, suggesting that the G300 implants have an increased ability to facilitate bone formation. A progression in the osteogenic cascade for implants spanning the medullary canal took place with residual inflammatory response, neovessel development along titanium struts, and some early formation of bone occurring inside and outside of scaffold pores after four weeks. By twelve weeks of implantation, bone formed in a somewhat unidirectional manner to span scaffold struts and infiltration of bone marrow occurred into the scaffold interior. Overall, the additively manufactured scaffolds both provide additional surface area for bone growth to occur inside the pores, a means to maintain the tissue through the vascular network, and a promising means for encouraging osseointegration at the implant interface.

## Acknowledgements

Work was performed at the McMaster Center for Advanced Light Microscopy, and McMaster’s Biointerfaces Institute. The authors would like to acknowledge funding from the Foshan Science and Technology Innovation Project (No. 2018IT100212), Natural Sciences and Engineering Research Council of Canada (RGPIN-2020-05722), and Canada Research Chairs Program. Funding support is also acknowledged from the Research Internship Abroad (BEPE) grant program #2022/04536-1 from the São Paulo Research Foundation (FAPESP). The authors would also like to acknowledge the scientific efforts of Bryan E.J. Lee and Chiara Micheletti toward the publication of this work. The graphical abstract was created using BioRender.com.

