## Supporting Figures 1-4 for "Osseointegration of functionally-graded Ti6Al4V porous implants: Histology of the pore network"

| Kathryn Grandfield | Mohamed A Elbestawi | Jianyu Chen |
|---|---|---|
| Email: | Email: | Email: |
| | | |

### Supporting Information

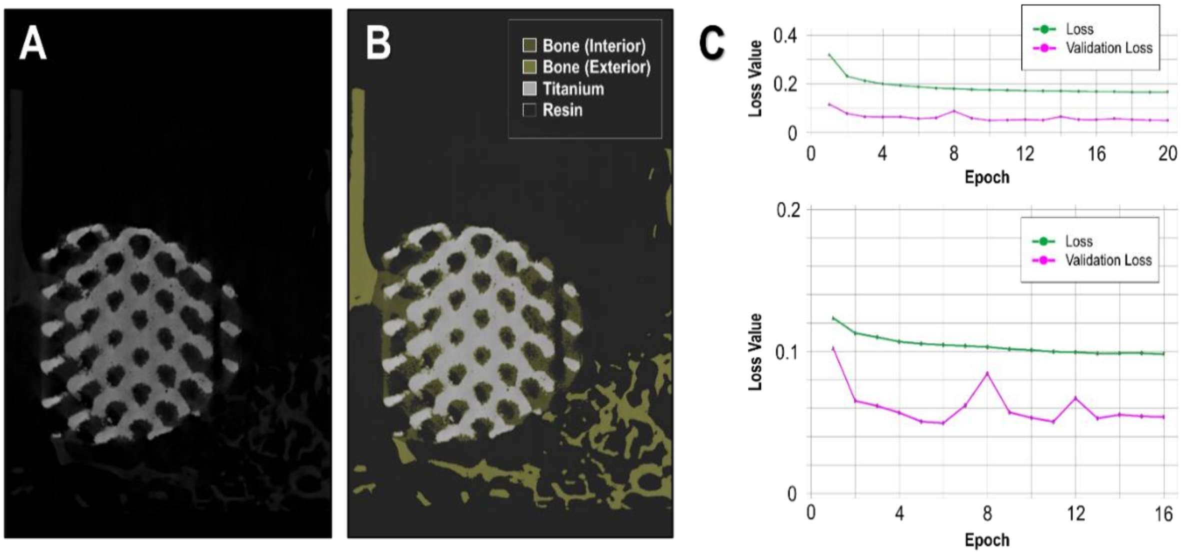


**Figure *S1: Convolutional neural network training on micro-CT data.*** *(A) Training slice from FG600 implant retrieved after four weeks. (B) Grayscale-assisted manual classification of bone inside the scaffold, bone outside the scaffold, titanium implant, and resin/void space for the slice presented in (A). (C) Training iterations of the convolutional neural network from ten slices of a FG600 scaffold and ten slices of a G300 scaffold. Loss values and validation loss values both decrease through successive epochs of training.*

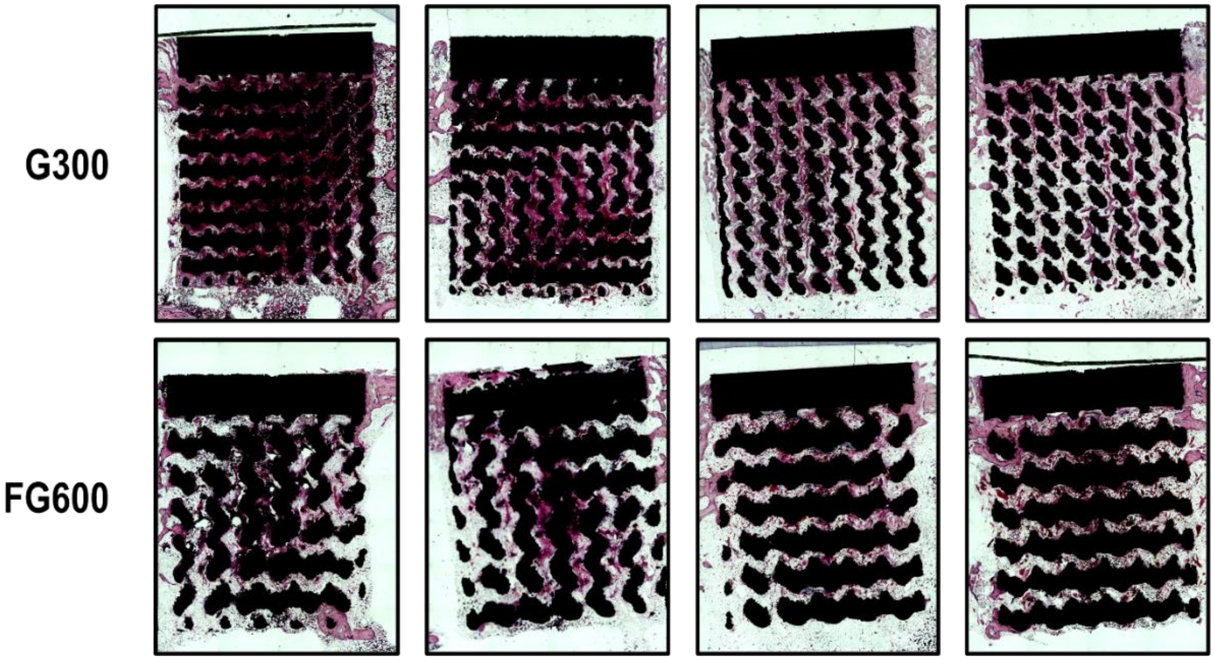


***Figure S2: All H&E stained sections from the four-week endpoint.*** *Implants in the same column were implanted bilaterally in the same rabbit, taking into account the effect of strong and weak responders. Common themes in all implants include inflammatory response occurring at the apex of the implant, cortical integration at the crown of the implant, and spindles of bone forming with microvasculature in the implant interior.*

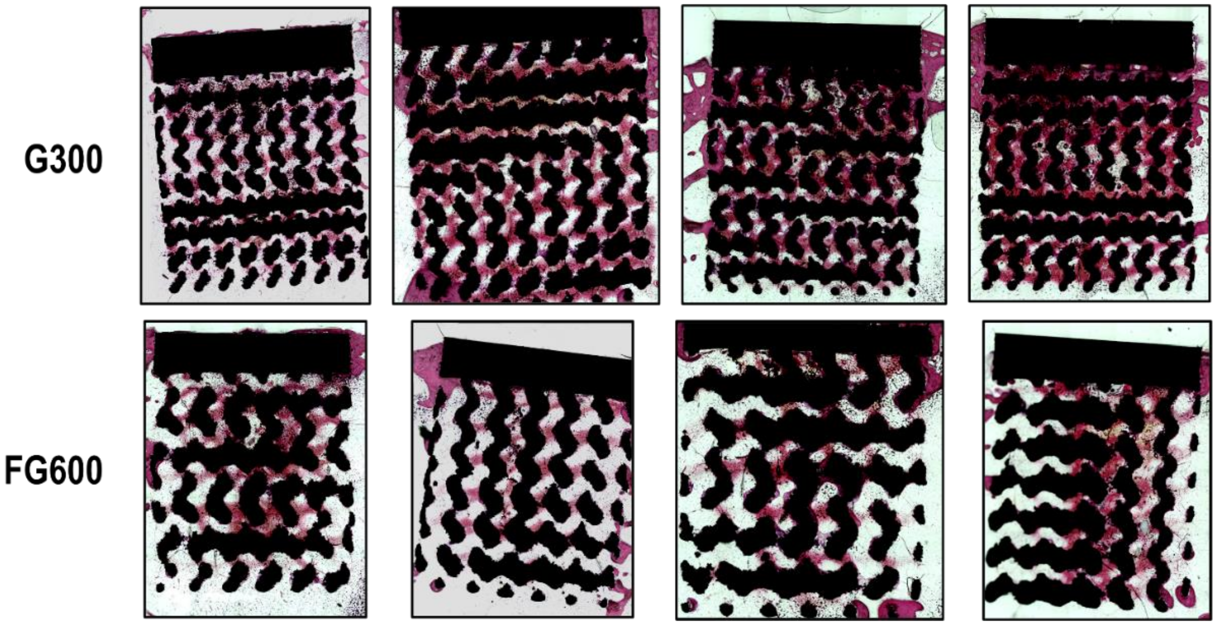


***Figure S3: All H&E stained sections from the twelve-week endpoint.*** *Implants in the same column were implanted bilaterally in the same rabbit, taking into account the effect of strong and weak responders. Bone apposition after twelve weeks appears more prolific in the G300 implants, and much of the inflammatory response has subsided. Regions of connective tissue bridge scaffold struts in the interior, and often appear unidirectional with tissue in neighbouring pore channels. Cortical and trabecular integration at the scaffold exterior appears coarser.*

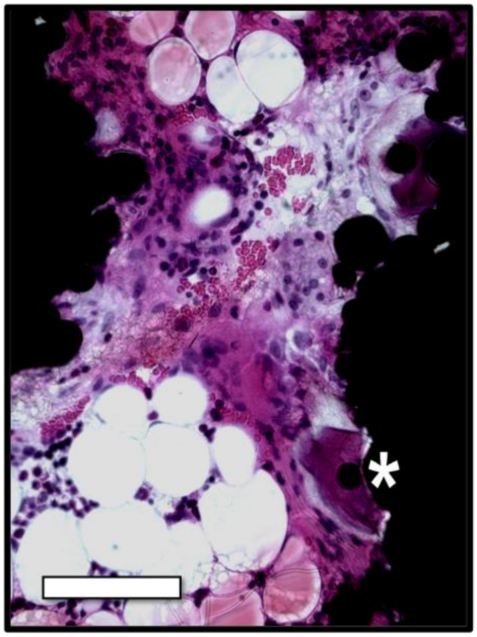

***Figure S4: Interior pore of a H&E stained section from a G300 implant after twelve weeks.*** *Red blood cells propagate in vasculature along the centerline of the pore, while a suspected megakaryocyte (denoted with an asterisk) produces platelets to preserve vascular response inside the scaffold. Scale bar: 100 µm.*
